# Electrochemical Evaluation of Ion Substituted-Hydroxyapatite on HeLa Cells Plasma Membrane Potential

**DOI:** 10.1101/440214

**Authors:** Bernard Owusu Asimeng, Elvis Kwason Tiburu, Elsie Effah Kuafmann, Lily Peamka, Claude Fiifi Hayford, Samuel Essien-Baidoo, Obed Korshie Dzikunu, Prince Atsu Anani

## Abstract

This study reports the electrochemical activities of a novel ion substituted-Hydroxyapatite material in contact with HeLa cells. The work was performed to evaluate the inhibitory effects of various concentrations of the material on the ion transfer mechanisms in HeLa cells. The materials (n=2: HAp1 and HAp3) were prepared at different stirring times from *Achatina achatina* snail shells and phosphate-containing solution. The structure of the materials and the trace elements concentration were evaluated using x-ray diffractometry and infrared spectrometry as well as atomic absorption spectroscopy. Electrochemical studies conducted on the cells, after 30 min of exposure to the materials, demonstrated differential responses as elucidated by cyclic voltammetry. The voltammograms revealed HAp1 to be non-redox whereas HAp3 was redox active. Minimal concentrations of HAp1 showed high anodic peak current when compared to the HeLa cells alone, indicating a hyperpolarization of the cells. The peak current gradually reduced as the concentration of HAp1 was increased, and then a sudden rise suggesting inhibition of the cell action potential. HAp3 showed a wavy pattern of the anodic peak current when the material concentration was varied. Peak currents of 0.92 and 0.57 nA were recorded for HAp1 and HAp3, respectively at the highest concentration of 5 *μ*L. The results suggest that different inhibitory mechanisms are at play on the voltage-gated ion channels of the cells, indicating the possibility of using the materials to achieve different cancer proliferation inhibition.

## 1. Introduction

Hydroxyapatite (HAp) is an inorganic material with chemical formula [Ca_10_(PO_4_)_6_(OH)_2_]. It is similar to the mineral component of the bone and has found much use in the tissue engineering industry owing to its biocompatible nature. It is used as a scaffold to promote bone cell migration, adhesion, and growth (1)(2). HAp possesses good adsorptive properties because of its negative and positive ion constituents. This property coupled with its biocompatibility, and controllable biodegradability has made it a preferred biomaterial for the delivery of cancer drugs to cancer sites (3). HAp nanoparticles, like other nanocarriers, respond to certain conditions like pH to release drugs in slow and metered doses at targeted sites (4). This reduces systemic toxicity and increases the solubility of the drug (5)(6), a solution to major limitations in chemotherapy. In spite of the advantages of nanoparticles in drug delivery, nanocarriers face design issues which include nanoparticle loading efficiency, as well as stability of nanoparticles with the attached ligands (7). Some investigations have thus been focused on using the nanoparticles as drugs and not just vehicles. Reports from various studies indicate that synthetic HAp nanoparticles, without any loaded drug, exhibit inhibitory properties on liver and lung cancer cells with lesser restraints on normal cells (8)(9)(10)(11). The cancer cells engulf the nanoparticles through endocytosis and the nanoparticles are then adsorbed onto ribosomes, interrupting the binding of mRNA. This is reported to disrupt the protein synthesis of the cells, thus inhibiting proliferation (8).

Chemotherapy, targeted drug delivery, and the use of the nanoparticles as described above, cause cancer cell death, thus preventing uncontrolled proliferation, through necrosis (forcing the cell to undergo premature death) (12). The cell membrane ruptures causing the organelles to stop functioning. This causes all the contents, including ions, in the cytoplasm to be released out of the cell. The ions then act as free radicals and pick other ions from the surrounding normal cells causing oxidative stresses (OS) which leads to cell lysis and inflammation of the surrounding tissues (13). Literature has reported of necrosis causing leukemia as a result of OS development from chemotherapy (14). In contrast to necrosis, apoptosis (programmed cell death) occurs routinely in normal cells during growth and aging in homeostatic mechanism to retain cell population in tissues (15). It is favorable since ions are not released into the extracellular space, thus avoiding the inflammatory response (16)(17). Apoptosis can be initiated via two pathways: intrinsic and extrinsic depending on the origin of the death-causing stimuli (18)(19). The pathways activate caspase (proteases that trigger the cell death process) which in turn requires reduction of potassium ions in the cytoplasm. Thus, ion flow across the plasma membrane and its subsequent depolarization (influx of positive ions in cell cytoplasm) are events that characterize apoptosis. Cell fragments known as apoptotic bodies are released after apoptosis. Microphages engulf the apoptotic bodies and remove them before the cell contents spill into the extracellular space (20).

It is reported that disrupting the plasma membrane potential of cancer cells may trigger apoptosis (21). Generally, cell plasma membranes generate potential (membrane potential) because of imbalance of charges between the intracellular and extracellular environment. This arises from the presence of different ion channels which allow distinct ions such as Na^+^, K^+^, Ca^2+^ and Cl^−^ access through the voltage-gates of the ion channels based on their specific size. Because of the difference in the number of ions within the cytoplasm and the extracellular medium, a voltage difference (electrochemical gradient) is always evolving. A normal cell tries to balance the ion concentration across the plasma membrane to achieve a resting potential through polarization. At resting potential, the cytoplasm becomes more negative. That is, more Na^+^ are sent outside and K^+^ are kept inside the cytoplasm (polarization). Typically, normal cells generate stimuli through polarization, for example, for the muscle to contract (22). On the contrary, cancer cells generate stimuli through plasma membrane depolarization (more Na^+^ are taken by the cell and the cytoplasm becomes less negative). Depolarization generates strong stimuli that communicate faster causing rapid proliferation (23).

To achieve apoptosis in cancer treatment, we have prepared a novel ion substituted-HAp material to influence the ion flow across the cell plasma membrane. The material is expected to release stimuli in the form of current when a low applied voltage is subjected to it. This will disrupt the depolarized membrane potential for apoptosis instigation. As proof of concept, the material is added to HeLa cells *in-vitro,* a potentiostat of a cyclic voltammetry (CV) supplies a potential to the mixture, and the ion transfer profiles from and to the cytoplasm recorded. The profiles are then related to the cell action potential.

## 2. Materials and Methods

### 2.1. Materials/Reagents

*Achatina achatina* (AA) snail shells were collected from snail sellers in Ghana and identified. Ammonium phosphate dibasic (APD) was purchased from Daejung chemicals & metals, Korea. HeLa cells (America Type Cell Culture (ATCC), USA) cultured in Roswell Park Memorial Institute (RPMI) medium 1640 (Sigma-Aldrich) supplemented with 10% Fetal Bovine Serum (FBS) and Penicillin-Streptomycin (PS: 100 U/mL) (Gibco, UK) were used in the cell studies.

### 2.2. HAp preparation

AA shells were calcined at 800°C using the method described by Asimeng et al (24). HAp materials (n=2) were prepared from 0.3 M of the calcined AA shell (CAAS) material, and 0.5 M APD solutions. 5.9 g of APD (pH 8.12) was added to 0.15 cm^3^ of distilled water to produce 0.5 M APD solution. 0.3 M CAAS solution was also produced by adding 5.7 g of CAAS powders to 0.15 cm^3^ of distilled water while stirring. The APD solution was added to the CAAS drop-wise and stirred using a magnetic stirrer hot plate for 1 and 3 hrs, respectively at a constant temperature of 40°C. These were labelled HAp1, and HAp3 for HAp stirred at 1 and 3 hrs, respectively. The materials were permitted to age for 24 hrs and then filtered. The filtrates were dried in an oven at a 100°C for 2 hrs to form HAp materials (24).

### 1.3. Characterization techniques

X-ray diffractometer (PANalytical Empyrean, Netherlands), with Cu Kα radiation, λ=1.5406 Å, was used to determine the phase compositions of the materials. The scan was in steps of 2° per min and the phases identified by comparing the material patterns to the Joint Committee on Powder Diffraction Standards (JCPDS) file. The crystallite size of HAp1 and HAp3 were determined using Debye-Scherer’s equation from the extreme peaks in the range of 30-35°. Infrared (IR) Spectrometer (Spectrum Two FT-IR, Perkin Elmer) was used to examine the functional groups present in the materials. The spectra were obtained over the region of 400-4000 cm^−1^ and compared to the Database of ATR-FT-IR spectra of various materials. Atomic Absorption Spectrometry (PinAAcle 900 AA Spectrometer, Perkin Elmer) of selectable spectral bandwidth 0.2, 0.7, and 2.0 nm was used to determine the concentration of trace elements in the materials.

### 2.4. Electrochemical and In-vitro cell studies with HAp materials

HeLa cells were cultured in T-75 culture flasks of RPMI-1640 supplemented with 10% FBS, 1% PS and kept in an incubator at 37°C and 5% CO2. HeLa cells were detached from culture flasks using trypsin and kept in suspension in culture medium. 5 mg of each HAp material was added to 50 *μ*L of a phosphate buffer saline (PBS) and microcentrifuged for 60 s after which varying concentrations (1-5 *μ*L) of the supernatant were pipetted into 100 *μ*L of HeLa cells. The bulk mixture was incubated for 30 min at room temperature before CV and viability studies using Cheap Stat Potentiostat (IORodeo, USA) with an Ag/AgCl interdigitated electrode (DropSens, UK), and a Cellometer Mini (Nexcelom, USA), respectively. 5 *μ*L of the bulk mixture for various concentrations of HAp1 and HAp3 were pipetted onto the electrode for CV readings. 20 *μ*L of the bulk mixture was added to 20 *μ*L of 0.2% trypan blue prepared in PBS (pH 7.2) after which 20 *μ*L was then pipetted on to the counting chamber of the cellometer for cell viability.

## 3. Results and Discussion

### 3.1. XRD analysis

Figure 1 shows XRD patterns for raw AA shell and CAAS materials. The raw powder material pattern in Figure 1(a) has aragonite structure. The aragonite was calcined at 800°C to convert into calcite, because calcite is more stable and commonly used for HAp preparation(25). The calcination temperature transforms aragonite to calcite and decomposes portions into calcium oxide (23). The pattern in Figure 1(b) shows the structure of the CAAS materials and this reveals phases of calcite and calcium hydroxide. Calcium oxide absorbs moisture from the atmosphere to form calcium hydroxide. Figure 2 shows the XRD patterns of HAp prepared from CAAS and APD. The patterns demonstrate a similar structure for HAp1 and HAp3. However, the variations in the stirring time resulted in different crystallite sizes. The crystallite size calculated for HAp1 and HAp3 were 40.86 ± 4.13 and 37.82 ± 3.48 nm, respectively.

**Figure 1:**
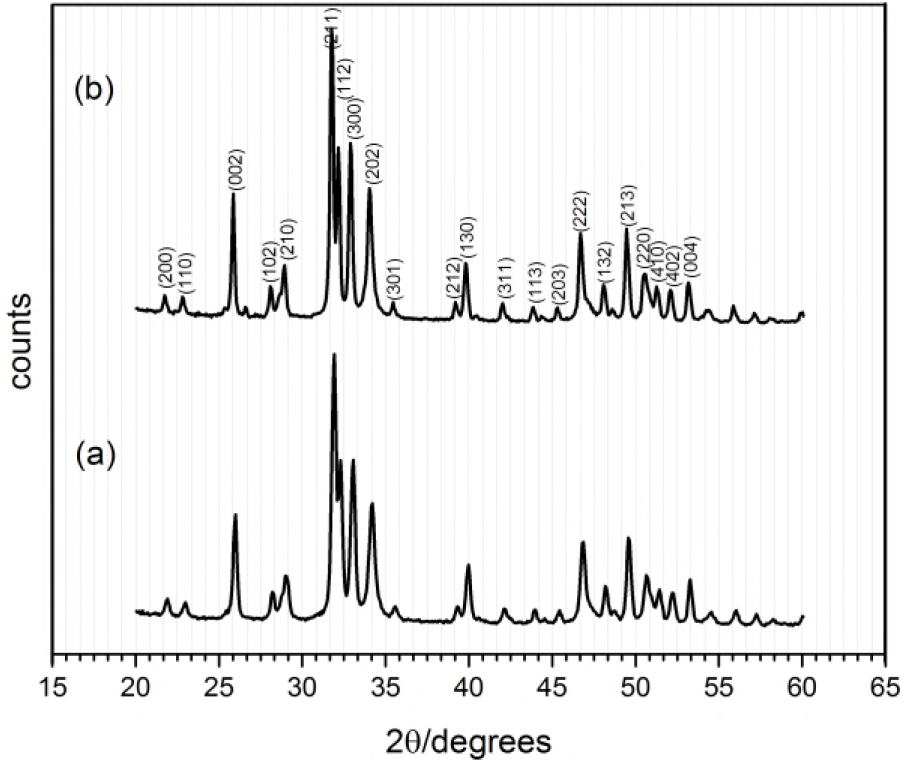
XRD patterns of HAp materials prepared at v tried stirring times (a) HAp1 (b) HAp3. The pattern looks similar with slight variation in the full half width maximum (FHWM).

**Figure 2:**
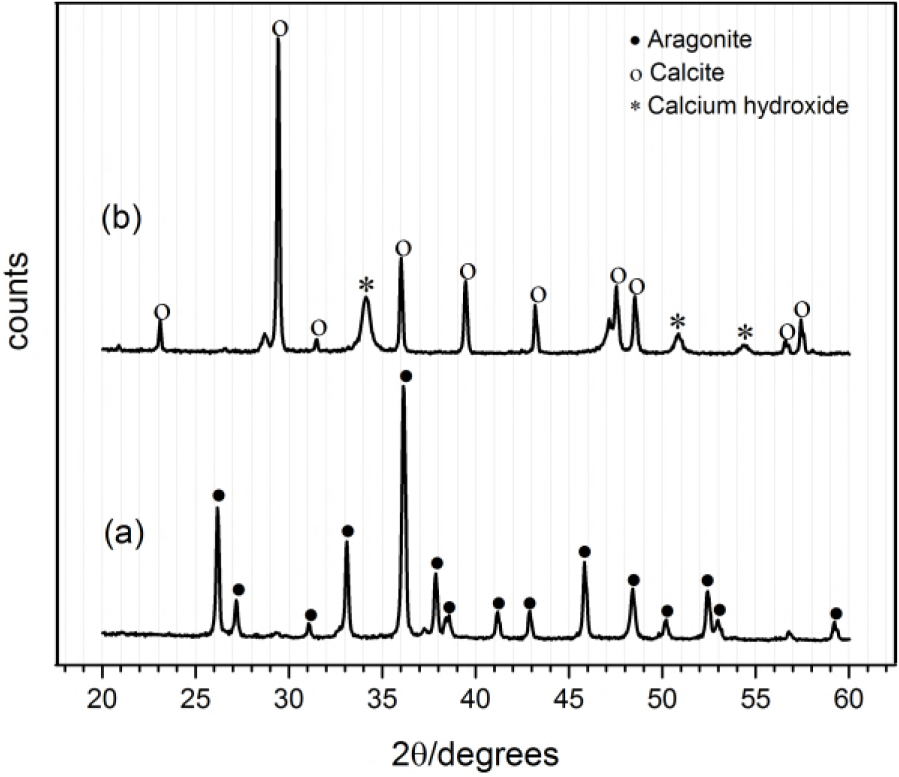
XRD patterns of AA shell materials (a) raw (b) CAAS at 800°C. The raw materials show pure phase of aragonite structure whereas the CAAS reveals major phases of calcite and minor phases of calcium hydroxide.

### 3.2. Infrared (IR) spectroscopy

FT-IR confirms the functional groups present in the prepared materials. The fingerprints in Figure 3(a) corresponded to the aragonite structure as confirmed from the Database of ATRFT-IR spectra of various materials (24)(26). The fingerprints with wavenumber 1082 cm^-1^ disappeared with CAAS. The fingerprints at wavenumbers 712, 874, and 1417 cm^−1^ in Figure 3(b) are those for calcite functional groups. These fingerprints support the XRD results of structural change from aragonite to calcite. The steep region around wavenumber 400 to 712 cm^−1^ indicates the presence of Ca-O stretching weak bonds (27) whereas the wavenumber at 3643 and 712 cm^−1^ are the fingerprints of calcium hydroxide (27). Figure 4 shows the fingerprints of HAp functional groups. The six phosphate groups occurred at wavenumbers 473, 563, 600, 963, 1029, and 1090 cm^−1^. This indicates the symmetric vibrational modes, *v*1 and *v*2 of phosphate at 473 and 963 cm^−1^, respectively whereas *v*3 appears at wavenumbers 1029 and 1090 cm^−1^. The final *v*4 shows at wavenumbers 563 and 600 cm^−1^. The two hydroxide groups of HAp appeared at wavenumbers 634 and 3574 cm^−1^. The carbonate groups at wavenumber 874 and 1412 cm^−1^ suggest that the materials are carbonated HAp. The carbonate peaks show a difference in the transmittance between HAp1 and HAp3. This variation is due to the prolonged stirring of the materials at 40°C as reported in literature (24). The decomposition of the carbonate might have been the reason for the reduction of the crystallite size indicated in the XRD.

**Figure 3:**
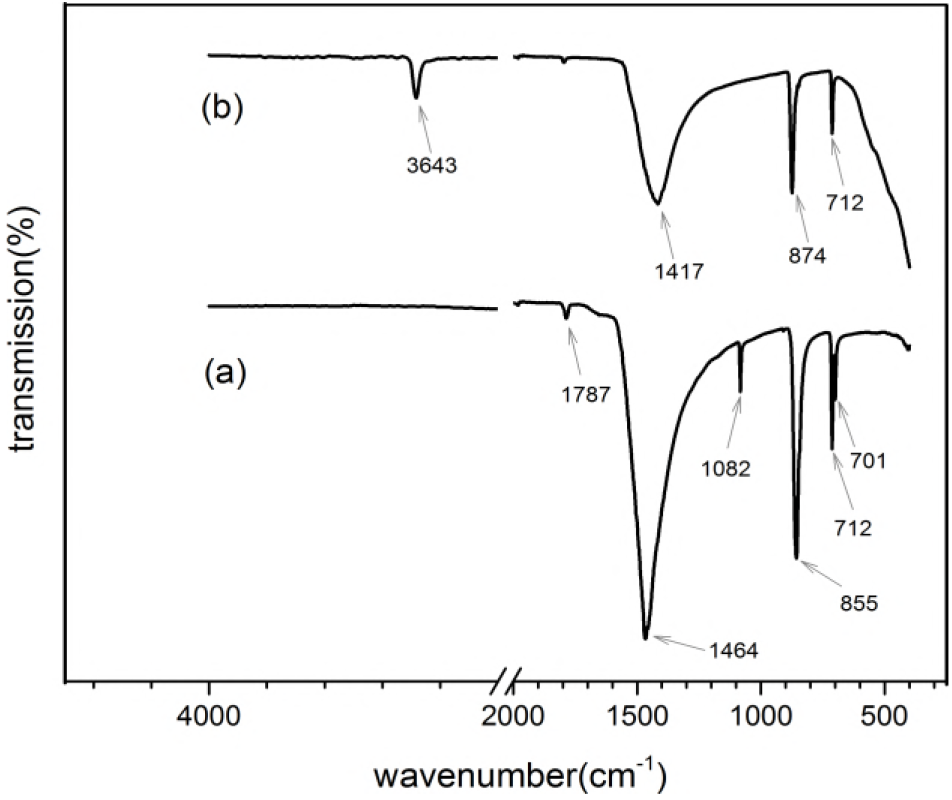
FT-IR spectra (a) raw (b)CAAS materials. The raw material corresponded to aragonite structure whereas the CAAS material is made up of calcite and hydroxide groups.

**Figure 4:**
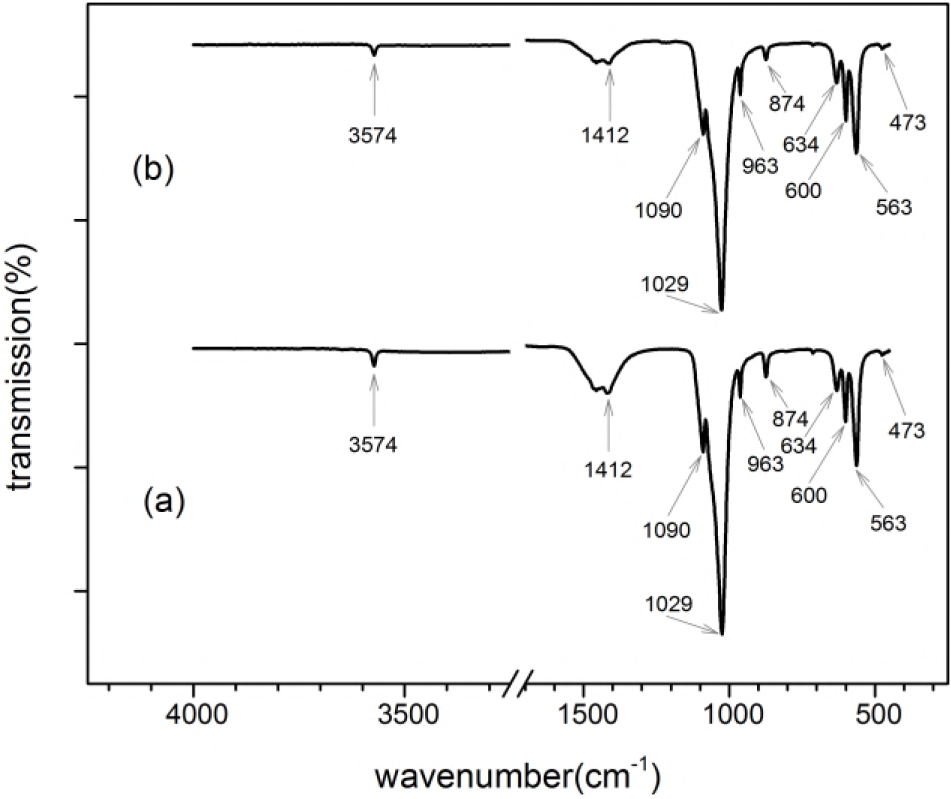
FT-IR spectra of the HAp materials prepared at varied stirring times (a) HAp1 (b) HAp3. The wavenumbers at 1412 and 874 cm^−1^ confirms the materials are carbonated-HAp. The differences in the carbonate finger prints were due to the prolonged stirring temperature.

### 3.3. AAS Studies

Figure 5 shows the concentrations of trace elements in HAp1 and HAp3. The AAS studies conducted on sodium (Na), potassium (K), zinc (Zn), magnesium (Mg), lead (Pb), and cadmium (Cd) reveals higher concentrations of Na, K, and Zn ions. Mg, Pb, and Cd ions were below the detection limit of the device. The bar graphs show a considerable difference in Na^+^ concentration in HAp1 and HAp3. The stirring times affected the concentration of Na^+^ that were available to the surface of the electrode of the potentiostat. HAp1 has fewer Na^+^ whereas HAp3 contains more surface ions, indicating ion substitutions in HAp1.

**Figure 5:**
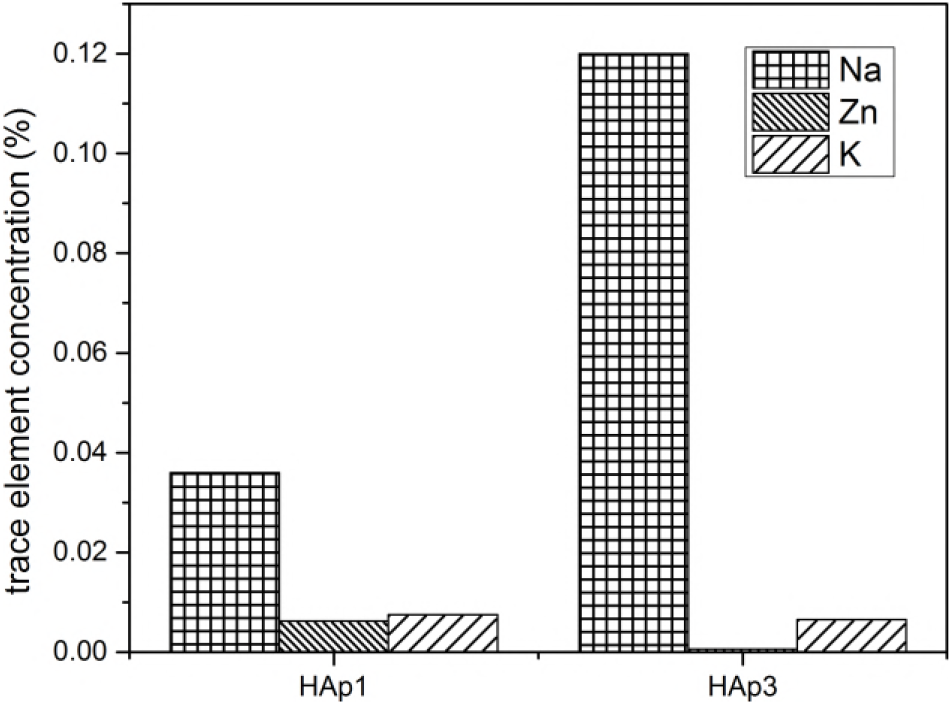
AAS results of HAp1 and HAp3. The materials have high Na ions than the other trace elements present. HAp1 has fewer surface ions than HAp3, indicating that more ions were substituted in H.

### 3.4. CV studies of cells with HAp materials

Figure 6 shows the cyclic voltammograms of cell medium alone, cell medium with HeLa (cmHeLa) cells, HAp1, and HAp3. The cell medium (RPMI-1460) contains inorganic salts, and amino acids and when placed on the working electrode of the potentiostat, exhibited redox activity. The cell medium alone gave electrons to the electrode at the oxidative half cycle and was recorded as an anodic peak current (545 μA). Conversely, when the potential sweep reversed, the electrode gave electrons back to the cell medium. This was recorded as a cathodic peak current on the reduction half cycle. The cmHeLa cells gave a higher anodic peak current than the cell medium alone. This result indicates that the cell released additional ions from the cytoplasm into the extracellular space (polarization), which is unusual with cancer cells. Normally, cancer cells generates depolarized membrane potential for the cell to proliferate (23). Therefore, the recorded behaviour, is due to the electrochemical gradient created as the electrode of the potentiostat supplied electrons to the medium, thus, making outside of the cytoplasm less positive. HAp1 and HAp3 activated in a PBS solution showed non-redox and redox activities, respectively. This trend supports the AAS results where HAp1 had fewer ions on its surface than HAp3. Figure 7 shows bar graphs of anodic peak currents against HAp concentrations. Addition of 1 μL of HAp1 into cmHeLa cells generated a higher anodic peak current compared to that of cmHeLa cells alone. It is suspected that substituted ions in the lattice sites of the HAp material adsorbed anions from the cell medium forming a layer. Literature reports indicate that a chemical called sialic acid released by the cell membrane of cancer cells promotes attachment of HAp to the cell surface, thus allowing much of the particles to be engulfed (28). It is hence suspected that, the cell attaches itself to the material and results in the formation of a double-layered cell when taken up; cell membrane of the HeLa cell as one layer and the internalized material as the other layer. The arrangement of the double layer together with the cytoplasmic materials and the electrode of the potentiostat is believed to behave as a capacitor and is evident as a spike in Figure 8 (A). That is, the spike represents a short period of direct current (Faradic current) which acts as a stimulus to hyperpolarize the cell during the oxidation half cycle. The cell’s voltage - gated ion channels opened, allowing more ions to move from the cytoplasm into the extracellular medium. Thus, the electrodes record a higher anodic peak current. The cytoplasm becomes more negative and restricts the proliferation signal that orders the cells to multiply. The cell tries to regain its optimum membrane potential by depolarizing as indicated in Figure 7. The anodic peak current decreases gradually as the concentration of HAp1 increases to 4 μL. The anodic peak current recorded at 4 μL is almost the same as that of the cell medium alone. This indicates that the concentrations of HAp1 from 2 to 4 μL did not generate stimuli to affect the cell’s ion flow across the membrane. As because as the concentration increased, the materials formed a stack structure (as illustrated with Scheme 1) which increased the double layer thickness, d, and in turn reducing the capacitance of the product formed, hence the material’s Faradic current. The anions in the medium are suspected to have been neutralized by the ions in the HAp1 material at 2 and 3 μL. However, Figure 8 reveals a negative spike, at the reduction half cycle of HAp1 at 4 μL, indicating an acceptance of electrons from the electrode to occupy free available sites. This caused a re-alignment of the dipoles formed, hence generating a Faradic current, B to activate a hyperpolarization at 5 μL. The higher anodic peak current recorded, indicates a higher number of ions that were transferred from the bulk mixture onto the electrode surface. This is suspected to cause cell rupture since depolarization of the plasma membrane is inhibited through the stimuli permanently opening the voltage-gates of the ion channel. Cell viability test on HAp1 at 5 μL using trypan blue diazo (a dye exclusion test) confirmed a low possibility of cells to survive (Figure 9). The dye exclusion test is based on the principle that damaged cells take up permeable dyes while viable cells are impermeable to the dye. Proteins from the damaged cells interact with the dye and fluoresce for detection by the cellometer, thus quantifying them as dead cells.

**Figure 6:**
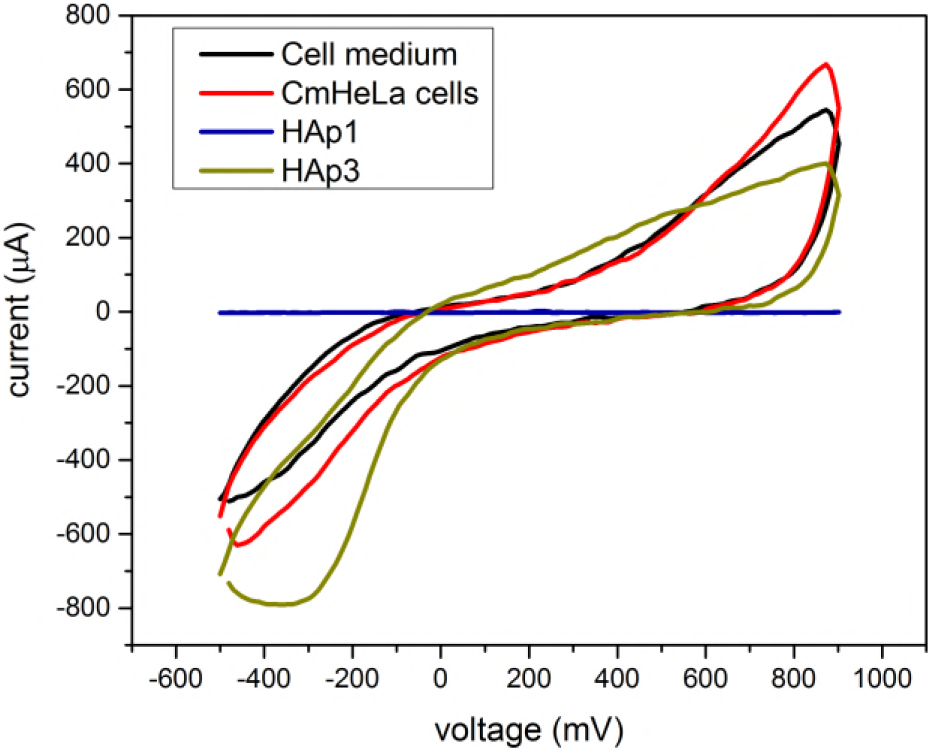
Cyclic voltammograms of cell medium, cell medium with HeLa (CmHeLa) cells, HAp1, and HAp3. All the materials were exhibiting redox activity except HAp1 alone which is non-redox active.

**Figure 7:**
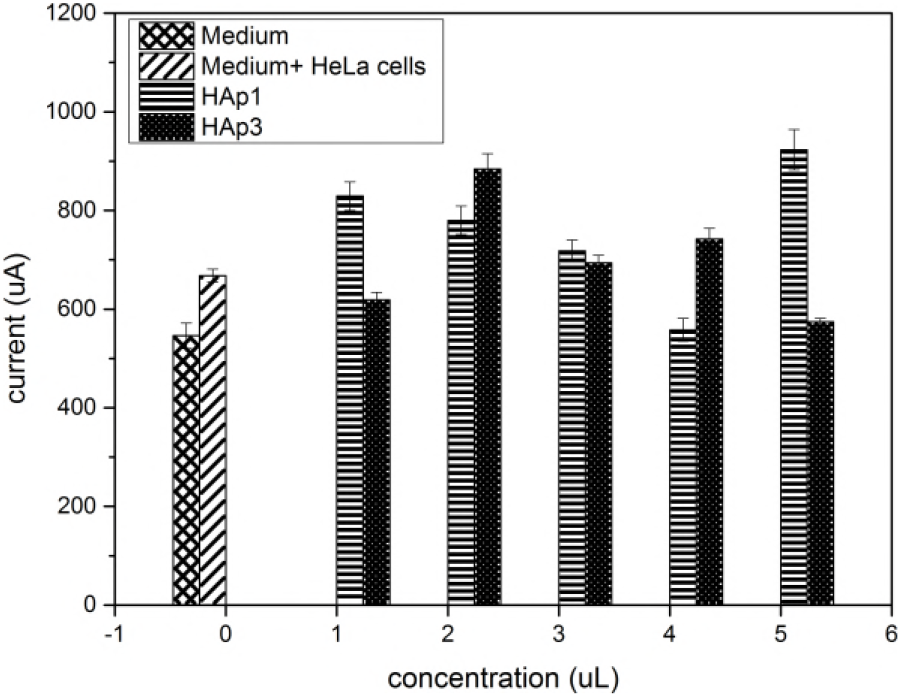
The influence of the HAp1, and HAp3 concentration on the anodic peak current of the HeLa cells.

**Figure 8:**
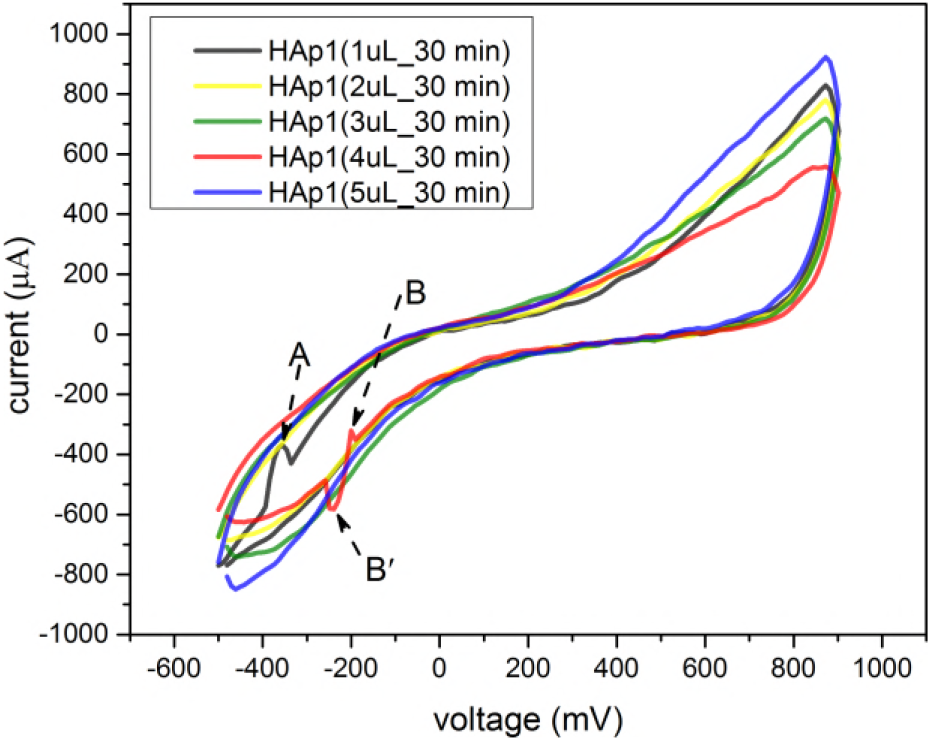
Cyclic voltammograms of CmHeLa cells with different concentrations of HAp1. A and B are oxidative peaks generated because of the double layer polarization whereas B’ is reductive peak indicating electron transfer from the electrode to the available free sites.

**Scheme 1:**
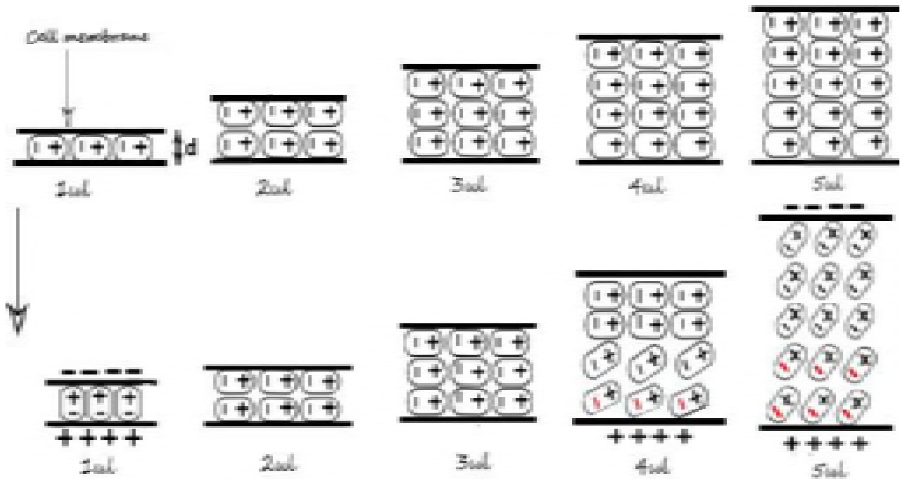
Proposed mechanistic behaviour of HAp1 on the HeLa cells. The cells form a double layer with the HAp1 material. Varying the concentration of HAp1 increases the layer thickness, d. The arrow indicates the behaviour of the layer when the potential is applied. The red signs are the electrons introduced by the electrode of the potentiostat during reduction half cycle.

**Figure 9:**
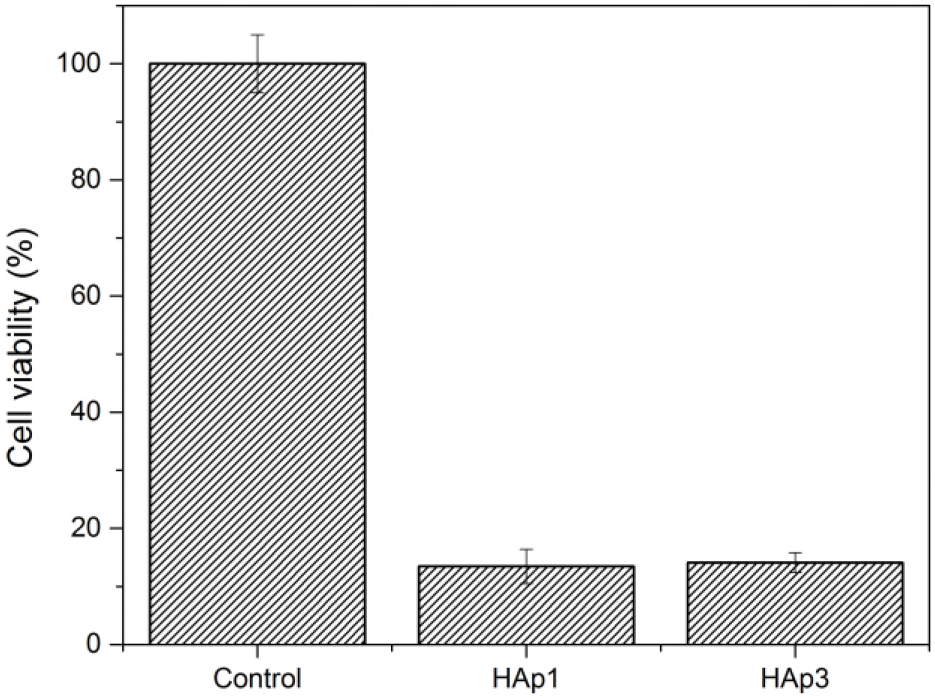
Cell viability studies of HAp1, and HAp3 materials of concentration, 5 μL each. Cell survival was low when the cells interacted with the HAp materials.

Addition of HAp3 to cmHeLa cells resulted in a wavy pattern indicating a different influence on the ion transfer of the bulk mixture from that of HAp1. From Figure 7, the addition of 1 μL HAp3 caused a depolarization, and the recorded anodic peak current became comparable to that of the cell medium alone. However, further increase in the concentration of HAp3 to 2 μL caused a hyperpolarization due to a high electrochemical gradient established by the presence of Na^+^ in the extracellular medium as illustrated in Scheme 2. The electrode recorded a higher efflux of the ions from the cells. As the concentration of the Na^+^ ions in the extracellular medium increases by an increase in the concentration of HAp3, the plasma membrane potential attempts to depolarize to sustain the cell’s integrity, but the voltage gates of the ion channels are suspected to be locked permanently at 5 μL. For the cell to survive, there is a suspicion of the opening of a new potassium channels for K^+^ efflux, which is evident in the reduction half of the CV (indicated as C in Figure 10). This supports literature reports, that altering the ion concentration in the cytoplasm is the main determinant of the survival of a cell (29). However, as shown in Figure 7, peak current recorded at 5 μL is the same as that recorded for the cell medium alone. This indicates a failure in membrane repolarization which according to reports, is associated with early stages of apoptosis (30). Cell viability test at 5 μL indicated low chances of cell survival.

**Scheme 2:**
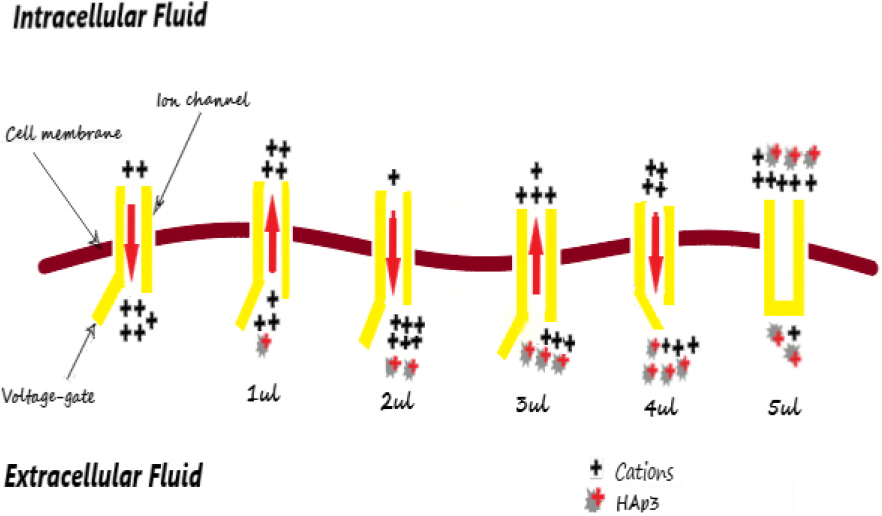
Proposed mechanistic behaviour of HAp3 on the HeLa cells. HAp3 adds up to the cation density in the extracellular fluid, creating a high potential gradient across the cell membrane to control the opening and closure of the voltage-gates.

**Figure 10:**
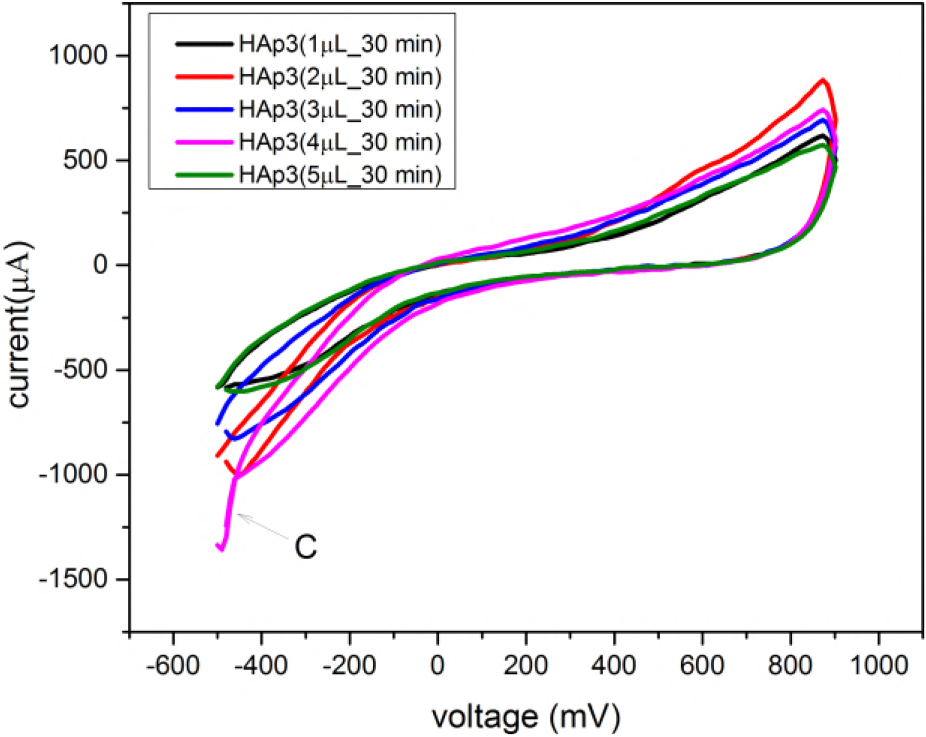
Cyclic voltammograms of CmHeLa cells with different concentrations of HAp3. C indicates the leakage of the Kpumps.

## 4. Conclusion

Electrochemical evaluation performed on ion substituted-HAp in HeLa cells reveals that the two materials prepared, HAp1 and HAp3, exhibit different ion transfer mechanisms. HAp1 and HAp3 are non-redox and redox active materials, respectively. The interaction of the HAp1 material with the cell medium and cells behave as a capacitor producing a Faradic current that controls ion flow across the cell plasma membrane. A Faradic current produced by the product at the reduction section, for higher concentrations of the material, is suspected to cause a cell rupture through permanent opening of the voltage-gates of the ion channels. On the other hand, HAp3 inhibited the voltage-gates of the ion channels through a permanent closure of the voltage-gate, therefore cell contents do not spill out unlike cell death achieved via necrosis. HAp1 and HAp3 have the potential to be used as low voltage assisted targeted anticancer materials and should be subjected to further biochemical studies to confirm that HAp3 is capable of inducing cancer cell death through apoptosis.

## Author Contributions

B. O. A conceived the idea, designed the experiment, and drafted the manuscript. E. K. T contributed analysis tool, laboratory resource and provided suggestions. E. E. K proofread the manuscripts and provided suggestions. L. P provided resources for the experiment and provided suggestions. C. F. H and S. E proofread and provided suggestions. 0.K.D and P. A. A were part of the data interpretation and manuscripts editing.

## Acknowledgements

Authors acknowledge A. K, T. Y. A. E, G. P. C. and E. A. A for their assistance during CV studies and cell culturing.

## Conflict of Interest

The authors have no conflict of interest to declare.

